## Supplemental figures for "A predicted structural interactome reveals binding interference from intrinsically disordered regions"

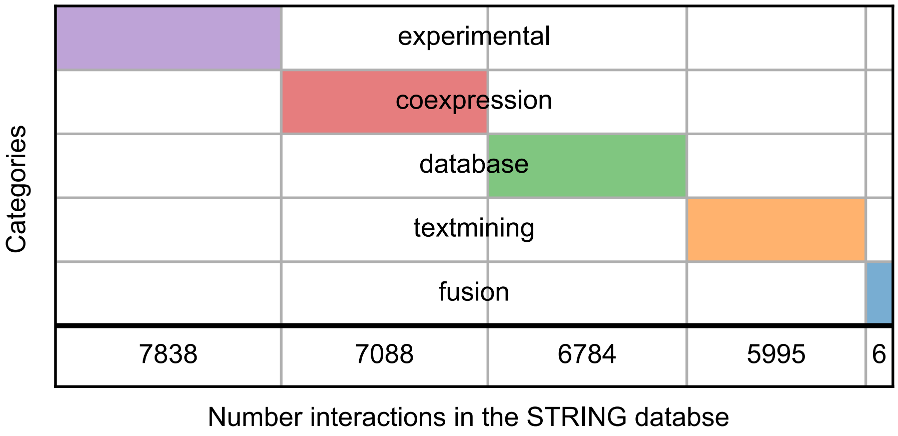


**S1 Fig. Number of predicted PPIs in different STRING database categories.**

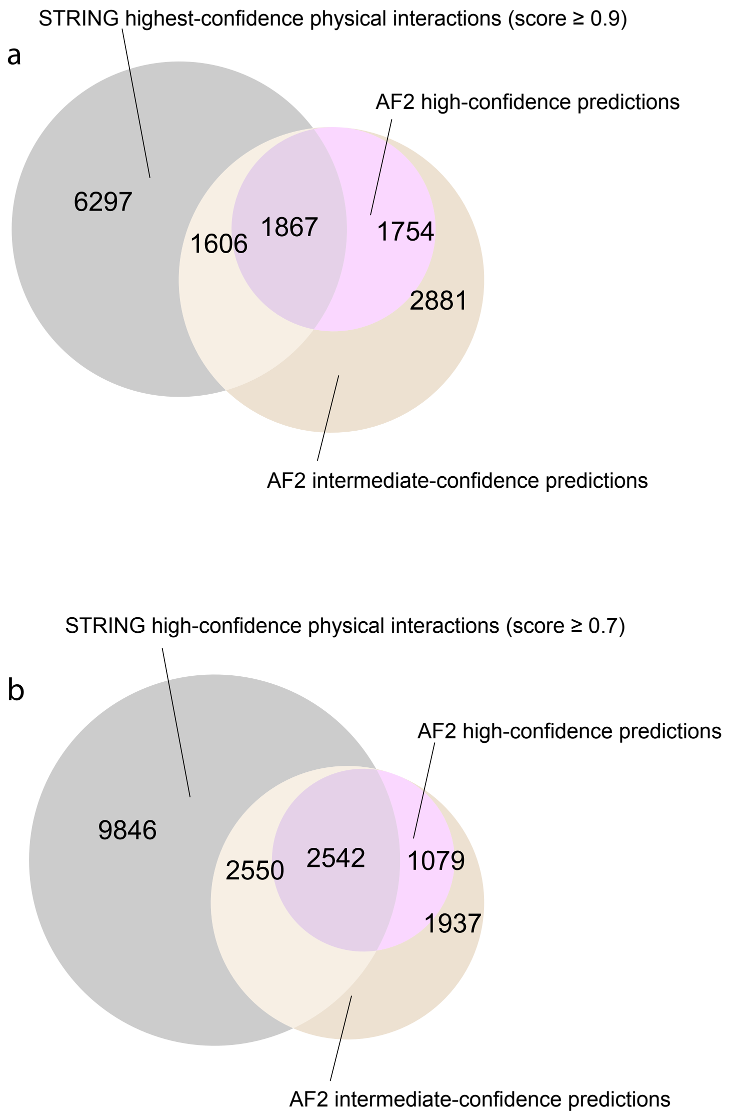


**S2 Fig. Overlap between high-confidence AlphaFold2 multimer predictions, intermediate-confidence AlphaFold2 multimer predictions, and the STRING database physical interactions**. A) The overlap between AlphaFold2 predictions and the highest-confidence STRING physical interactions (STRING physical confidence score ≥ 0.9). B) The overlap between AlphaFold2 predictions and the high-confidence STRING physical interactions (STRING physical confidence score ≥ 0.7).

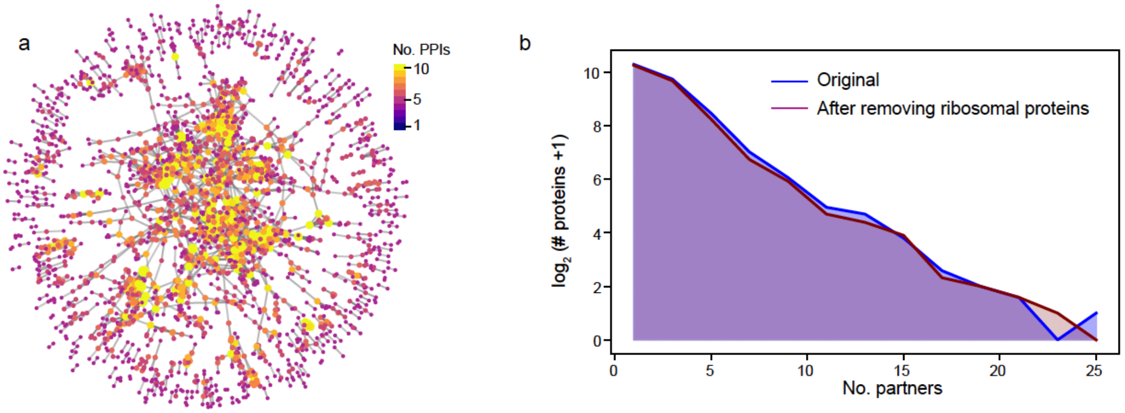


**S3 Fig. High-confidence PPI network after removing ribosomal proteins.** A) The modified network, obtained after removing 154 ribosomal proteins, including both cytoplasmic and mitochondrial, retains a similar overall layout to the original network (Fig 1B in the main text). B) The modified network also shows a comparable connectivity distribution to the original network. The connectivity distribution represents the number of proteins that have a given number of interaction partners. Note that the Y-axis is in log scale, $\log_2$(Number of proteins + 1).

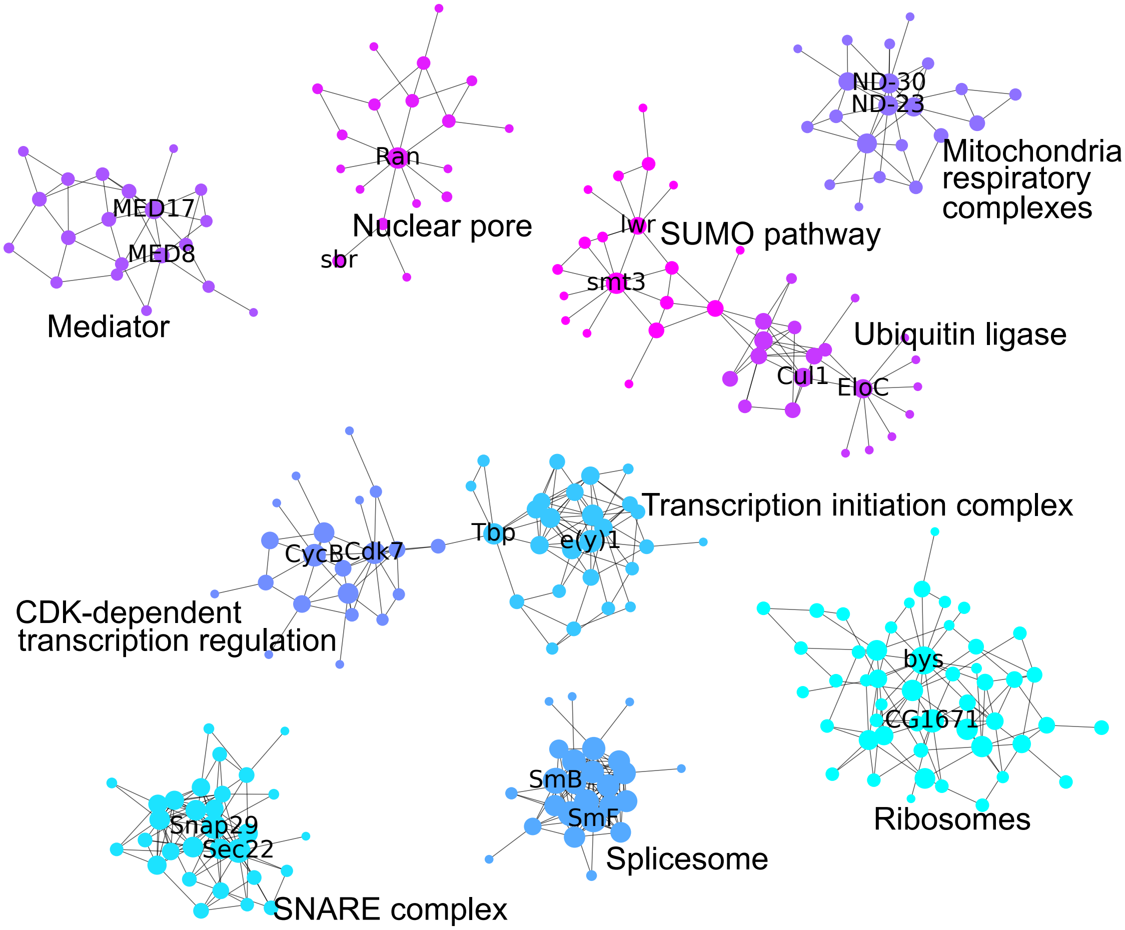


**S4 Fig. Top 10 MCL clusters of predicted high confidence PPIs**. Each cluster corresponds to an essential protein complex or a crucial cellular pathway.

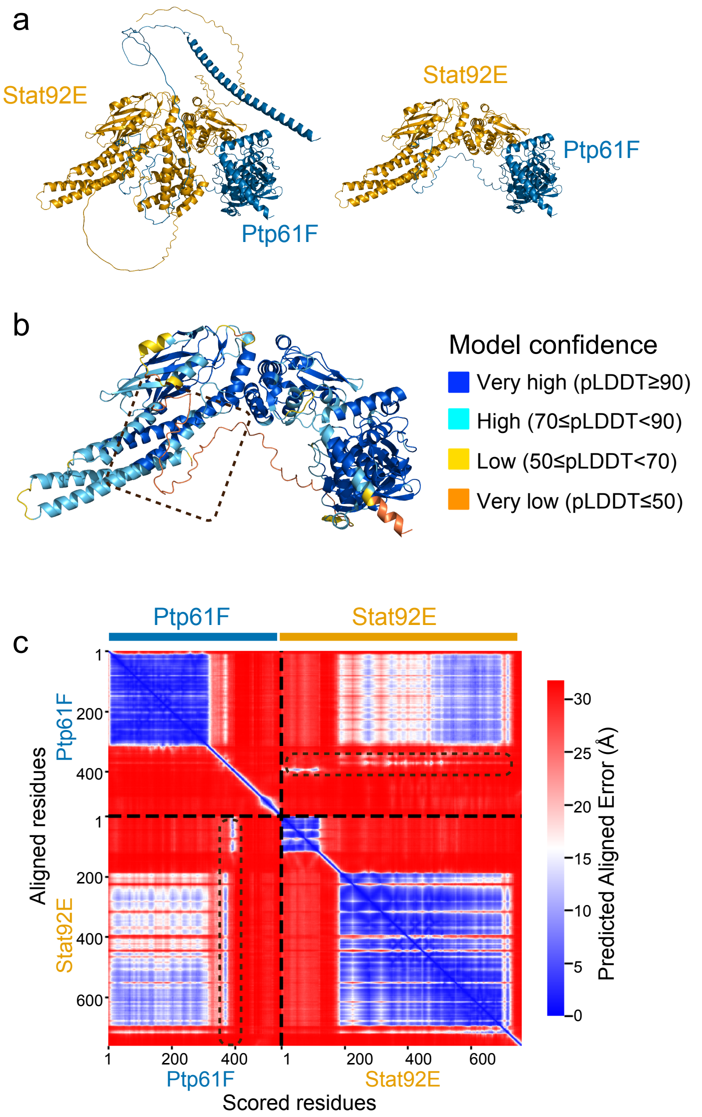


**S5 Fig. Prediction details of the Ptp61F–Stat92E interaction**. A) Left: full-length predicted model. Right: truncated model with the C-terminus of Ptp61F (residues 380–548), the N-terminus of Stat92E (residues 1–187), and the C-terminus of Stat92E (residues 696–961) removed for clearer visualization. B) Model confidence (pLDDT) for the truncated model shown in (A). The dashed box highlights a region where binding involves a low-confidence segment of Ptp61F (residues 360–370). C) Predicted Aligned Error (PAE) matrix. The blue off-diagonal region highlighted by the dashed box corresponds to the potential binding region indicated in (B).

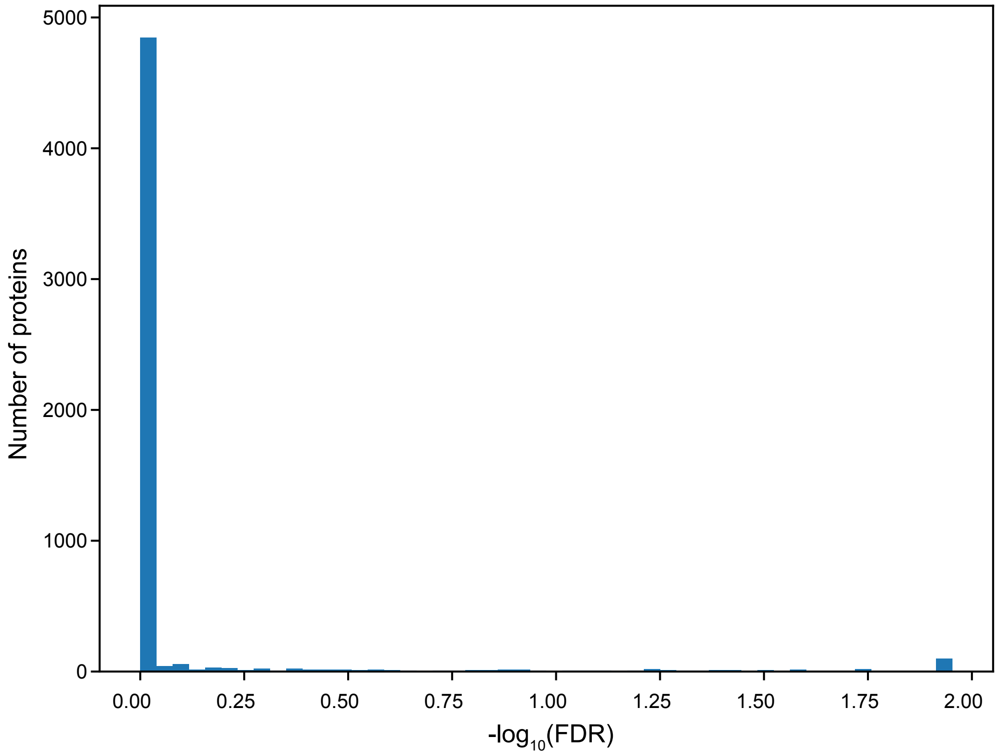


**S6 Fig. Permutation analysis indicates random distribution of prediction failures.** In our analysis, a total of 34,940 predictions were attempted, of which 27,711 were successfully predicted and 7,229 failed due to memory limitations. To determine whether prediction failures were randomly distributed across proteins, we performed a permutation test in which success labels were randomly reassigned 5,000 times while maintaining the total number of successful predictions. For each protein, empirical p-values were computed from the permutation distribution of success fractions and corrected for multiple testing using the Benjamini-Hochberg false discovery rate (FDR) method. The resulting FDR distribution shows that most proteins have FDR values close to 1 and thus $-\log_{10}$(FDR)s close to 0, with only a small number of outliers, suggesting that prediction failures were largely random rather than systematically associated with particular proteins or interaction types.

**S1 Table. Functional enrichment analysis of the largest connected subnetwork of predicted high-confidence PPIs**. Only GO terms with false discover rate smaller than 1e-50 were included.

| GO-term | Description | Count in Network | Strength | False Discovery Rate |
|---|---|---|---|---|
| **GO:0009987** | Cellular process | 1011 of 3942 | 0.39 | 6.89E-149 |
| **GO:0044237** | Cellular metabolic process | 896 of 3352 | 0.63 | 2.77E-136 |
| **GO:0043170** | Macromolecule metabolic process | 510 of 1090 | 0.45 | 1.27E-132 |
| **GO:0010467** | Gene expression | 704 of 2292 | 0.49 | 8.97E-130 |
| **GO:0044260** | Cellular macromolecule metabolic process | 617 of 1804 | 0.33 | 1.92E-127 |
| **GO:0034641** | Cellular nitrogen compound metabolic process | 907 of 3813 | 0.28 | 5.83E-125 |
| **GO:0006807** | Nitrogen compound metabolic process | 1035 of 4906 | 0.32 | 2.51E-116 |
| **GO:0008152** | Metabolic process | 930 of 4070 | 0.29 | 2.05E-98 |
| **GO:0044238** | Primary metabolic process | 953 of 4398 | 0.55 | 1.17E-85 |
| **GO:0071704** | Organic substance metabolic process | 425 of 1095 | 0.48 | 8.09E-85 |
| **GO:0090304** | Nucleic acid metabolic process | 452 of 1373 | 0.43 | 2.14E-84 |
| **GO:0006139** | Nucleobase-containing compound metabolic process | 510 of 1723 | 0.6 | 1.08E-83 |
| **GO:0044267** | Cellular protein metabolic process | 332 of 763 | 0.46 | 1.27E-81 |
| **GO:0034645** | Cellular macromolecule biosynthetic process | 462 of 1454 | 0.45 | 6.44E-81 |
| **GO:0046483** | Heterocycle metabolic process | 465 of 1497 | 0.33 | 3.83E-79 |
| **GO:0006725** | Cellular aromatic compound metabolic process | 673 of 2859 | 0.44 | 1.82E-77 |
| **GO:0071840** | Cellular component organization or biogenesis | 469 of 1552 | 0.58 | 2.9E-75 |
| **GO:1901360** | Organic cyclic compound metabolic process | 320 of 767 | 0.55 | 1.33E-61 |
| **GO:0044271** | Cellular nitrogen compound biosynthetic process | 331 of 841 | 0.44 | 1.93E-59 |
| **GO:0016070** | RNA metabolic process | 380 of 1257 | 0.43 | 1.62E-58 |
| **GO:0044249** | Cellular biosynthetic process | 373 of 1245 | 0.73 | 2.91E-58 |
| **GO:0044085** | Cellular component biogenesis | 182 of 310 | 0.42 | 7.03E-58 |
| **GO:0006412** | Translation | 382 of 1314 | 0.42 | 2.64E-57 |
| **GO:0009058** | Biosynthetic process | 377 of 1290 | 0.6 | 1.26E-56 |
| **GO:1901576** | Organic substance biosynthetic process | 234 of 540 | 0.33 | 5.3E-56 |
| **GO:0006396** | RNA processing | 522 of 2235 | 0.73 | 5.54E-54 |

**S2 Table. Functional enrichment analysis of the second largest connected subnetwork of predicted high-confidence PPIs**.

| GO-term | Description | Count in Network | Strength | False Discovery Rate |
|---|---|---|---|---|
| **GO:0032504** | Multicellular organism reproduction | 13 of 1226 | 0.66 | 0.00030 |
| **GO:0000003** | Reproduction | 14 of 1361 | 0.65 | 0.00014 |
| **GO:0050794** | Regulation of cellular process | 19 of 3805 | 0.34 | 0.0286 |

**S3 Table. Functional enrichment analysis of the third largest connected subnetwork of predicted high-confidence PPIs.** Only biological process with false discovery rate smaller than 1e-10 were displayed.

| GO-term | Description | Count in Network | Strength | False Discovery Rate |
|---|---|---|---|---|
| **GO:0007178** | Transmembrane receptor protein serine/threonine kinase signaling | 12 of 18 | 2.63 | 3.02E-24 |
| **GO:0060395** | SMAD protein signal transduction | 10 of 14 | 2.66 | 5.05E-20 |
| **GO:0030509** | BMP signaling pathway | 10 of 15 | 2.63 | 5.61E-20 |
| **GO:0071363** | Cellular response to growth factor stimulus | 11 of 44 | 2.2 | 8.66E-19 |
| **GO:0007476** | Imaginal disc-derived wing morphogenesis | 12 of 264 | 1.46 | 1.05E-12 |
| **GO:0071310** | Cellular response to organic substance | 12 of 282 | 1.43 | 1.85E-12 |
| **GO:0045464** | R8 cell fate specification | 7 of 20 | 2.35 | 3.76E-12 |
| **GO:0032924** | Activin receptor signaling pathway | 6 of 8 | 2.68 | 6.72E-12 |
| **GO:0090092** | Regulation of transmembrane receptor protein serine/threonine signaling | 8 of 54 | 1.97 | 8.54E-12 |
| **GO:0048639** | Positive regulation of developmental growth | 9 of 101 | 1.75 | 1.08E-11 |
| **GO:0010033** | Response to organic substance | 13 of 492 | 1.22 | 1.44E-11 |
| **GO:0050803** | Regulation of synapse structure or activity | 10 of 192 | 1.52 | 3.48E-11 |

**S4 Table. List of the 15 genomes to access the origin of IDR regions.**

| Species | Genome version |
|---|---|
| ***D.melanogaster*** | STRING database version 11.5 |
| ***D.simulans*** | GCF_016746395.2_Prin_Dsim_3.1 |
| ***D.yakuba*** | GCF_016746365.2_Prin_Dyak_Tai18E2_2.1 |
| ***D.erecta*** | GCF_003286155.1_DereRS2 |
| ***D.ananassae*** | GCF_017639315.1_ASM1763931v2 |
| ***D.willistoni*** | GCF_018902025.1_UCI_dwil_1.1 |
| ***D.grimshawi*** | GCF_018153295.1_ASM1815329v1 |
| ***S.lebanonensis*** | GCF_003285725.1_SlebRS2 |
| ***A.gambiae*** | GCF_943734735.2_idAnoGambNW_F1_1 |
| ***T.castaneum*** | GCF_031307605.1_icTriCast1.1 |
| ***D.magna*** | GCF_020631705.1_ASM2063170v1.1 |
| ***C.elegans*** | GCF_000002985.6_WBcel235 |
| ***D.rerio*** | GCF_000002035.6_GRCz11 |
| ***S.cerevisiae*** | S288C reference genome release 64-4-1 |
| ***E.coli*** | GCF_000005845.2_ASM584v2 |
| ***H.salinarum*** | GCF_004799605.1_ASM479960v1 |
