## Supplementary material for "A predicted structural interactome reveals binding interference from intrinsically disordered regions": S2 Appendix

cluster    1 1534 2630FBpp0071046,FBpp0293061,FBpp0087118,FBpp0087094,FBpp0079182,FBpp0072145,FBpp0081234,FBpp0301847,FBpp0073445,FBpp0087461,FBpp0083893,FBpp0072674,FBpp0071449,FBpp0088015,FBpp0080488,FBpp0071822,FBpp0076756,FBpp0074237,FBpp0075684,FBpp0079148,FBpp0074151,FBpp0073943,FBpp0070723,FBpp0075686,FBpp0079306,FBpp0099642,FBpp0071596,FBpp0078984,FBpp0073328,FBpp0077159,FBpp0079946,FBpp0074180,FBpp0303680,FBpp0077401,FBpp0087343,FBpp0087731,FBpp0077022,FBpp0070719,FBpp0304730,FBpp0071288,FBpp0110435,FBpp0072144,FBpp0070279,FBpp0271782,FBpp0075807,FBpp0072503,FBpp0085379,FBpp0083891,FBpp0079641,FBpp0289298,FBpp0100176,FBpp0079536,FBpp0080389,FBpp0071142,FBpp0081916,FBpp0082645,FBpp0071128,FBpp0082062,FBpp0072975,FBpp0072468,FBpp0305832,FBpp0079538,FBpp0087921,FBpp0305095,FBpp0304248,FBpp0085724,FBpp0303878,FBpp0070654,FBpp0088416,FBpp0083801,FBpp0083572,FBpp0082928,FBpp0070382,FBpp0071553,FBpp0078689,FBpp0111818,FBpp0292260,FBpp0078448,FBpp0071808,FBpp0079514,FBpp0075620,FBpp0083581,FBpp0082459,FBpp0075676,FBpp0078929,FBpp0303869,FBpp0082153,FBpp0087203,FBpp0079943,FBpp0073120,FBpp0073558,FBpp0075302,FBpp0073035,FBpp0071198,FBpp0080872,FBpp0087335,FBpp0074288,FBpp0072446,FBpp0291433,FBpp0070409,FBpp0083687,FBpp0084806,FBpp0087583,FBpp0079443,FBpp0078278,FBpp0082984,FBpp0071685,FBpp0302793,FBpp0290840,FBpp0083376,FBpp0070730,FBpp0100187,FBpp0082511,FBpp0077111,FBpp0070561,FBpp0089041,FBpp0305943,FBpp0087553,FBpp0076371,FBpp0113081,FBpp0070471,FBpp0305165,FBpp0087821,FBpp0083413,FBpp0074129,FBpp0078024,FBpp0100175,FBpp0071137,FBpp0076207,FBpp0087569,FBpp0079500,FBpp0086066,FBpp0301731,FBpp0077735,FBpp0099686,FBpp0088021,FBpp0081957,FBpp0083415,FBpp0072720,FBpp0100182,FBpp0085489,FBpp0087935,FBpp0070299,FBpp0088355,FBpp0077353,FBpp0071543,FBpp0073989,FBpp0072239,FBpp0300874,FBpp0087511,FBpp0081117,FBpp0083659,FBpp0087775,FBpp0081879,FBpp0072052,FBpp0087359,FBpp0085717,FBpp0084761,FBpp0071193,FBpp0072020,FBpp0082298,FBpp0081324,FBpp0086314,FBpp0079780,FBpp0305141,FBpp0082353,FBpp0087189,FBpp0079328,FBpp0070175,FBpp0078093,FBpp0088040,FBpp0087323,FBpp0071846,FBpp0084729,FBpp0080121,FBpp0082973,FBpp0074758,FBpp0074227,FBpp0074841,FBpp0071381,FBpp0079060,FBpp0074284,FBpp0071049,FBpp0084082,FBpp0082957,FBpp0291369,FBpp0304449,FBpp0086249,FBpp0086507,FBpp0077213,FBpp0086603,FBpp0086749,FBpp0084739,FBpp0073316,FBpp0077141,FBpp0085193,FBpp0083665,FBpp0078997,FBpp0071052,FBpp0085100,FBpp0084907,FBpp0074609,FBpp0099583,FBpp0079542,FBpp0083658,FBpp0078592,FBpp0079445,FBpp0080894,FBpp0074709,FBpp0300652,FBpp0083769,FBpp0081600,FBpp0074246,FBpp0072957,FBpp0297285,FBpp0073144,FBpp0076460,FBpp0079609,FBpp0072801,FBpp0070646,FBpp0305742,FBpp0075839,FBpp0083263,FBpp0075119,FBpp0073524,FBpp0086103,FBpp0082329,FBpp0088519,FBpp0071497,FBpp0110439,FBpp0083843,FBpp0081384,FBpp0080391,FBpp0070885,FBpp0087115,FBpp0074558,FBpp0086711,FBpp0083962,FBpp0083802,FBpp0077473,FBpp0071958,FBpp0074229,FBpp0078664,FBpp0080402,FBpp0079164,FBpp0087897,FBpp0084067,FBpp0089084,FBpp0080401,FBpp0087340,FBpp0081786,FBpp0080045,FBpp0084617,FBpp0305695,FBpp0087278,FBpp0079312,FBpp0080641,FBpp0078992,FBpp0080412,FBpp0082011,FBpp0073777,FBpp0075124,FBpp0088818,FBpp0112426,FBpp0078918,FBpp0075068,FBpp0072041,FBpp0073354,FBpp0305426,FBpp0082767,FBpp0081082,FBpp0071023,FBpp0079305,FBpp0081822,FBpp0078066,FBpp0078007,FBpp0072475,FBpp0081725,FBpp0072081,FBpp0081845,FBpp0080319,FBpp0084050,FBpp0110433,FBpp0085891,FBpp0087498,FBpp0305729,FBpp0084341,FBpp0070766,FBpp0291479,FBpp0110080,FBpp0288698,FBpp0077927,FBpp0080102,FBpp0082682,FBpp0076359,FBpp0078633,FBpp0076549,FBpp0077778,FBpp0077714,FBpp0083932,FBpp0086676,FBpp0305561,FBpp0086812,FBpp0304269,FBpp0084766,FBpp0085119,FBpp0087045,FBpp0305611,FBpp0071518,FBpp0082392,FBpp0079677,FBpp0302771,FBpp0082232,FBpp0078315,FBpp0077415,FBpp0304059,FBpp0083076,FBpp0077885,FBpp0070402,FBpp0076848,FBpp0074075,FBpp0084242,FBpp0074984,FBpp0085546,FBpp0077230,FBpp0088044,FBpp0291495,FBpp0070859,FBpp0071087,FBpp0078205,FBpp0074662,FBpp0071879,FBpp0302563,FBpp0082513,FBpp0084103,FBpp0076200,FBpp0087350,FBpp0079812,FBpp0080062,FBpp0073016,FBpp0078978,FBpp0071248,FBpp0100177,FBpp0080256,FBpp0080564,FBpp0304831,FBpp0085902,FBpp0078754,FBpp0077965,FBpp0072097,FBpp0078055,FBpp0099776,FBpp0079262,FBpp0082574,FBpp0078203,FBpp0087637,FBpp0304214,FBpp0081584,FBpp0100180,FBpp0077974,FBpp0305302,FBpp0078508,FBpp0303832,FBpp0074496,FBpp0073310,FBpp0080211,FBpp0289083,FBpp0084561,FBpp0071766,FBpp0078400,FBpp0080120,FBpp0113035,FBpp0297299,FBpp0082893,FBpp0072557,FBpp0075034,FBpp0080052,FBpp0082445,FBpp0305723,FBpp0082065,FBpp0083860,FBpp0305946,FBpp0075982,FBpp0073267,FBpp0071503,FBpp0085658,FBpp0112017,FBpp0072187,FBpp0297142,FBpp0078847,FBpp0087342,FBpp0289788,FBpp0076602,FBpp0075633,FBpp0304467,FBpp0085254,FBpp0078667,FBpp0078831,FBpp0075833,FBpp0306232,FBpp0303780,FBpp0073762,FBpp0072119,FBpp0077073,FBpp0074239,FBpp0302752,FBpp0071753,FBpp0077705,FBpp0304503,FBpp0075700,FBpp0290998,FBpp0271901,FBpp0076134,FBpp0305676,FBpp0075690,FBpp0076152,FBpp0087865,FBpp0076723,FBpp0300615,FBpp0297292,FBpp0089182,FBpp0076330,FBpp0076804,FBpp0076457,FBpp0070652,FBpp0075104,FBpp0083569,FBpp0290815,FBpp0084210,FBpp0078896,FBpp0073719,FBpp0084157,FBpp0080803,FBpp0087709,FBpp0082737,FBpp0074554,FBpp0084616,FBpp0071194,FBpp0076782,FBpp0082817,FBpp0083036,FBpp0087676,FBpp0111294,FBpp0079072,FBpp0077650,FBpp0088360,FBpp0081459,FBpp0076789,FBpp0305673,FBpp0291653,FBpp0070911,FBpp0305541,FBpp0072687,FBpp0071045,FBpp0073623,FBpp0100183,FBpp0079484,FBpp0074690,FBpp0078694,FBpp0082591,FBpp0086096,FBpp0084786,FBpp0071993,FBpp0085569,FBpp0303728,FBpp0079031,FBpp0075069,FBpp0072615,FBpp0073673,FBpp0070953,FBpp0085482,FBpp0293948,FBpp0075274,FBpp0305289,FBpp0081548,FBpp0076918,FBpp0082477,FBpp0074715,FBpp0072659,FBpp0304981,FBpp0112455,FBpp0305446,FBpp0303867,FBpp0082998,FBpp0078568,FBpp0083975,FBpp0088016,FBpp0087648,FBpp0302559,FBpp0084721,FBpp0071283,FBpp0294012,FBpp0087004,FBpp0304812,FBpp0077396,FBpp0081545,FBpp0071232,FBpp0070814,FBpp0085716,FBpp0076990,FBpp0086348,FBpp0090943,FBpp0294035,FBpp0073681,FBpp0070304,FBpp0086519,FBpp0083436,FBpp0076827,FBpp0082570,FBpp0081040,FBpp0087475,FBpp0070143,FBpp0083857,FBpp0289620,FBpp0099655,FBpp0070877,FBpp0078473,FBpp0083570,FBpp0086013,FBpp0074641,FBpp0079470,FBpp0081263,FBpp0071252,FBpp0073090,FBpp0072150,FBpp0073651,FBpp0111299,FBpp0070637,FBpp0087500,FBpp0088810,FBpp0074835,FBpp0298005,FBpp0076738,FBpp0078688,FBpp0078894,FBpp0088035,FBpp0100185,FBpp0081001,FBpp0304879,FBpp0080997,FBpp0070679,FBpp0080578,FBpp0079873,FBpp0082433,FBpp0071848,FBpp0304829,FBpp0070284,FBpp0080484,FBpp0084623,FBpp0084735,FBpp0076424,FBpp0084345,FBpp0084778,FBpp0088489,FBpp0087863,FBpp0297500,FBpp0081860,FBpp0087883,FBpp0076464,FBpp0070259,FBpp0271716,FBpp0072083,FBpp0081820,FBpp0081343,FBpp0071102,FBpp0070805,FBpp0079785,FBpp0084894,FBpp0081385,FBpp0078532,FBpp0112244,FBpp0086705,FBpp0076352,FBpp0072233,FBpp0072184,FBpp0082743,FBpp0078095,FBpp0082469,FBpp0085888,FBpp0081305,FBpp0073459,FBpp0082462,FBpp0070316,FBpp0083753,FBpp0086115,FBpp0070302,FBpp0304876,FBpp0080827,FBpp0304068,FBpp0079399,FBpp0297441,FBpp0070950,FBpp0074058,FBpp0082134,FBpp0080747,FBpp0082867,FBpp0112985,FBpp0305828,FBpp0086872,FBpp0081029,FBpp0297199,FBpp0082833,FBpp0070363,FBpp0078316,FBpp0070716,FBpp0288680,FBpp0083371,FBpp0081488,FBpp0302532,FBpp0088275,FBpp0297239,FBpp0081599,FBpp0302981,FBpp0084899,FBpp0083434,FBpp0070370,FBpp0084648,FBpp0086387,FBpp0082137,FBpp0088833,FBpp0078416,FBpp0083126,FBpp0087596,FBpp0084875,FBpp0113099,FBpp0087651,FBpp0073036,FBpp0083502,FBpp0072876,FBpp0071794,FBpp0080441,FBpp0082877,FBpp0074687,FBpp0071303,FBpp0070618,FBpp0076819,FBpp0083625,FBpp0077909,FBpp0291726,FBpp0085104,FBpp0303726,FBpp0077106,FBpp0083195,FBpp0074565,FBpp0070650,FBpp0077679,FBpp0075618,FBpp0083740,FBpp0078990,FBpp0076264,FBpp0081544,FBpp0084626,FBpp0291070,FBpp0084936,FBpp0078583,FBpp0070940,FBpp0112484,FBpp0083319,FBpp0291670,FBpp0080629,FBpp0087566,FBpp0077946,FBpp0077099,FBpp0080775,FBpp0081756,FBpp0074584,FBpp0086400,FBpp0292263,FBpp0304487,FBpp0087322,FBpp0082970,FBpp0072103,FBpp0074664,FBpp0077079,FBpp0083779,FBpp0077516,FBpp0075010,FBpp0070150,FBpp0076645,FBpp0081493,FBpp0071587,FBpp0073230,FBpp0293859,FBpp0076374,FBpp0304751,FBpp0079565,FBpp0077209,FBpp0077125,FBpp0305182,FBpp0081592,FBpp0075317,FBpp0083015,FBpp0076458,FBpp0074717,FBpp0082314,FBpp0070430,FBpp0099945,FBpp0076339,FBpp0072154,FBpp0080967,FBpp0099511,FBpp0086098,FBpp0083892,FBpp0081412,FBpp0076791,FBpp0305117,FBpp0086531,FBpp0301016,FBpp0070351,FBpp0302537,FBpp0074481,FBpp0301842,FBpp0076672,FBpp0099891,FBpp0078662,FBpp0086732,FBpp0086881,FBpp0072494,FBpp0071994,FBpp0070141,FBpp0079628,FBpp0289288,FBpp0073336,FBpp0082995,FBpp0078415,FBpp0305700,FBpp0293269,FBpp0073149,FBpp0084971,FBpp0081004,FBpp0077142,FBpp0077694,FBpp0081216,FBpp0079616,FBpp0084955,FBpp0074887,FBpp0300957,FBpp0085589,FBpp0301099,FBpp0074847,FBpp0071771,FBpp0080870,FBpp0075801,FBpp0077206,FBpp0078961,FBpp0100186,FBpp0086223,FBpp0083131,FBpp0072990,FBpp0083089,FBpp0305335,FBpp0086599,FBpp0070298,FBpp0070976,FBpp0080701,FBpp0072745,FBpp0292865,FBpp0302530,FBpp0290461,FBpp0110412,FBpp0077378,FBpp0088345,FBpp0073018,FBpp0079418,FBpp0078348,FBpp0073963,FBpp0077309,FBpp0304615,FBpp0100181,FBpp0086294,FBpp0076028,FBpp0297920,FBpp0072134,FBpp0074500,FBpp0077630,FBpp0083564,FBpp0289558,FBpp0081380,FBpp0084893,FBpp0086616,FBpp0081333,FBpp0293589,FBpp0070415,FBpp0082467,FBpp0083989,FBpp0111841,FBpp0080706,FBpp0084787,FBpp0290353,FBpp0080560,FBpp0071970,FBpp0076558,FBpp0293261,FBpp0290878,FBpp0297417,FBpp0304862,FBpp0077107,FBpp0077900,FBpp0085121,FBpp0072132,FBpp0086063,FBpp0088018,FBpp0072130,FBpp0076849,FBpp0087085,FBpp0072768,FBpp0070808,FBpp0110561,FBpp0291442,FBpp0303920,FBpp0271847,FBpp0077859,FBpp0084375,FBpp0294011,FBpp0073083,FBpp0303766,FBpp0072425,FBpp0078167,FBpp0088375,FBpp0070403,FBpp0080051,FBpp0084486,FBpp0076433,FBpp0078970,FBpp0088995,FBpp0083105,FBpp0070793,FBpp0088483,FBpp0086402,FBpp0113022,FBpp0074115,FBpp0086953,FBpp0078856,FBpp0297300,FBpp0083646,FBpp0084348,FBpp0087347,FBpp0112014,FBpp0112147,FBpp0085308,FBpp0077790,FBpp0072179,FBpp0075701,FBpp0075718,FBpp0073058,FBpp0074529,FBpp0081076,FBpp0087712,FBpp0070709,FBpp0078317,FBpp0080867,FBpp0079592,FBpp0110413,FBpp0074650,FBpp0078919,FBpp0305286,FBpp0071034,FBpp0084754,FBpp0086965,FBpp0111848,FBpp0083912,FBpp0080143,FBpp0084918,FBpp0087484,FBpp0078053,FBpp0084351,FBpp0081856,FBpp0079892,FBpp0271779,FBpp0085563,FBpp0071675,FBpp0084342,FBpp0078516,FBpp0082522,FBpp0074122,FBpp0086012,FBpp0075626,FBpp0306023,FBpp0080808,FBpp0086578,FBpp0082177,FBpp0084574,FBpp0086269,FBpp0303329,FBpp0078758,FBpp0070242,FBpp0084244,FBpp0291394,FBpp0073099,FBpp0081124,FBpp0081848,FBpp0079769,FBpp0087645,FBpp0070862,FBpp0304603,FBpp0082158,FBpp0079444,FBpp0293514,FBpp0084150,FBpp0080552,FBpp0290365,FBpp0078296,FBpp0081322,FBpp0082592,FBpp0077151,FBpp0084966,FBpp0081397,FBpp0086901,FBpp0292527,FBpp0292384,FBpp0080905,FBpp0084831,FBpp0073411,FBpp0089035,FBpp0078760,FBpp0300193,FBpp0292305,FBpp0304887,FBpp0084959,FBpp0070907,FBpp0085616,FBpp0289999,FBpp0083168,FBpp0086776,FBpp0072946,FBpp0086101,FBpp0075704,FBpp0079833,FBpp0303669,FBpp0074665,FBpp0081435,FBpp0072249,FBpp0087861,FBpp0075382,FBpp0080647,FBpp0076142,FBpp0077998,FBpp0073781,FBpp0086469,FBpp0079332,FBpp0075754,FBpp0082468,FBpp0074822,FBpp0088562,FBpp0083516,FBpp0077259,FBpp0305661,FBpp0075166,FBpp0074660,FBpp0072998,FBpp0077500,FBpp0086874,FBpp0081281,FBpp0087936,FBpp0081156,FBpp0077316,FBpp0303792,FBpp0110418,FBpp0082568,FBpp0306039,FBpp0072388,FBpp0072037,FBpp0070638,FBpp0111355,FBpp0070349,FBpp0081350,FBpp0072660,FBpp0112092,FBpp0085566,FBpp0077551,FBpp0079880,FBpp0304312,FBpp0074855,FBpp0084713,FBpp0070355,FBpp0290897,FBpp0079052,FBpp0074128,FBpp0304481,FBpp0085893,FBpp0079680,FBpp0077749,FBpp0290542,FBpp0290035,FBpp0080670,FBpp0305904,FBpp0087765,FBpp0304202,FBpp0082018,FBpp0077775,FBpp0083206,FBpp0297918,FBpp0083273,FBpp0079295,FBpp0071651,FBpp0073004,FBpp0071600,FBpp0087707,FBpp0073822,FBpp0293232,FBpp0083884,FBpp0075952,FBpp0085915,FBpp0071222,FBpp0072602,FBpp0075713,FBpp0086107,FBpp0080890,FBpp0291583,FBpp0078433,FBpp0074971,FBpp0082540,FBpp0080146,FBpp0293863,FBpp0081673,FBpp0081650,FBpp0099795,FBpp0082071,FBpp0100184,FBpp0290208,FBpp0304594,FBpp0071259,FBpp0072561,FBpp0305498,FBpp0271702,FBpp0086206,FBpp0081351,FBpp0085829,FBpp0075947,FBpp0079607,FBpp0084059,FBpp0113107,FBpp0083451,FBpp0083134,FBpp0083022,FBpp0078681,FBpp0306049,FBpp0087663,FBpp0080255,FBpp0073983,FBpp0302048,FBpp0076342,FBpp0078624,FBpp0070007,FBpp0077625,FBpp0080329,FBpp0084763,FBpp0290742,FBpp0081807,FBpp0070624,FBpp0079874,FBpp0081491,FBpp0081560,FBpp0082735,FBpp0084638,FBpp0087272,FBpp0072061,FBpp0110163,FBpp0293882,FBpp0084051,FBpp0072312,FBpp0075663,FBpp0076695,FBpp0085258,FBpp0079417,FBpp0081665,FBpp0085166,FBpp0083780,FBpp0070864,FBpp0085585,FBpp0079527,FBpp0073913,FBpp0077720,FBpp0072342,FBpp0085678,FBpp0072209,FBpp0078483,FBpp0073490,FBpp0077658,FBpp0087216,FBpp0303776,FBpp0076831,FBpp0083927,FBpp0077783,FBpp0081606,FBpp0292155,FBpp0072113,FBpp0078673,FBpp0086505,FBpp0081355,FBpp0072976,FBpp0081317,FBpp0301081,FBpp0083348,FBpp0087519,FBpp0081834,FBpp0072598,FBpp0081683,FBpp0110594,FBpp0087536,FBpp0073828,FBpp0084094,FBpp0305616,FBpp0074906,FBpp0083976,FBpp0070479,FBpp0088121,FBpp0076312,FBpp0081900,FBpp0082849,FBpp0073290,FBpp0071563,FBpp0084933,FBpp0304617,FBpp0302789,FBpp0071703,FBpp0297168,FBpp0297577,FBpp0070114,FBpp0070729,FBpp0070093,FBpp0305699,FBpp0110252,FBpp0070860,FBpp0303146,FBpp0304140,FBpp0088069,FBpp0081703,FBpp0074214,FBpp0077618,FBpp0088654,FBpp0076798,FBpp0083354,FBpp0085295,FBpp0099564,FBpp0073608,FBpp0071028,FBpp0305521,FBpp0306036,FBpp0305462,FBpp0085013,FBpp0072151,FBpp0289334,FBpp0077644,FBpp0085480,FBpp0077277,FBpp0079772,FBpp0073380,FBpp0077445,FBpp0089217,FBpp0082590,FBpp0075997,FBpp0112228,FBpp0081484,FBpp0086853,FBpp0072906,FBpp0070954,FBpp0076859,FBpp0074801,FBpp0080998,FBpp0305371,FBpp0086597,FBpp0290169,FBpp0073493,FBpp0305864,FBpp0290503,FBpp0070203,FBpp0076005,FBpp0271789,FBpp0072050,FBpp0084784,FBpp0085801,FBpp0078500,FBpp0099829,FBpp0076232,FBpp0075277,FBpp0080490,FBpp0305422,FBpp0079537,FBpp0085539,FBpp0288575,FBpp0079278,FBpp0077582,FBpp0082387,FBpp0300877,FBpp0288961,FBpp0084989,FBpp0304226,FBpp0086850,FBpp0305840,FBpp0075969,FBpp0085954,FBpp0086285,FBpp0074818,FBpp0081614,FBpp0073386,FBpp0078993,FBpp0075761,FBpp0305877,FBpp0111476,FBpp0081894,FBpp0085222,FBpp0074184,FBpp0079776,FBpp0303852,FBpp0083958,FBpp0072803,FBpp0072065,FBpp0082803,FBpp0084078,FBpp0076777,FBpp0305911,FBpp0303635,FBpp0301175,FBpp0080880,FBpp0071213,FBpp0290883,FBpp0072893,FBpp0085479,FBpp0085530,FBpp0071541,FBpp0089396,FBpp0087063,FBpp0289644,FBpp0077936,FBpp0076607,FBpp0070764,FBpp0088441,FBpp0077899,FBpp0085192,FBpp0070398,FBpp0079818,FBpp0293540,FBpp0080622,FBpp0079266,FBpp0071455,FBpp0081295,FBpp0072055,FBpp0074054,FBpp0080039,FBpp0071277,FBpp0070913,FBpp0078949,FBpp0112403,FBpp0306150,FBpp0082569,FBpp0082667,FBpp0080799,FBpp0289256,FBpp0078375,FBpp0085517,FBpp0111983,FBpp0074607,FBpp0099759,FBpp0077357,FBpp0112136,FBpp0306028,FBpp0080879,FBpp0305266,FBpp0076372,FBpp0078901,FBpp0075901,FBpp0078382,FBpp0080395,FBpp0290792,FBpp0086767,FBpp0088945,FBpp0304189,FBpp0082547,FBpp0076832,FBpp0077263,FBpp0074514,FBpp0076244,FBpp0075937,FBpp0078844,FBpp0086135,FBpp0085334,FBpp0074686,FBpp0112051,FBpp0075938,FBpp0082384,FBpp0071459,FBpp0073513,FBpp0079054,FBpp0082731,FBpp0085224,FBpp0087491,FBpp0082110,FBpp0070358,FBpp0074528,FBpp0301993,FBpp0304246,FBpp0082888,FBpp0072908,FBpp0082231,FBpp0072056,FBpp0082285,FBpp0070045,FBpp0072142,FBpp0111894,FBpp0072657,FBpp0075617,FBpp0076601,FBpp0076418,FBpp0112453,FBpp0086010,FBpp0078353,FBpp0078032,FBpp0071064,FBpp0078765,FBpp0082219,FBpp0072463,FBpp0077495,FBpp0271844,FBpp0085744,FBpp0072658,FBpp0079409,FBpp0083075,FBpp0112122,FBpp0079468,FBpp0078891,FBpp0076917,FBpp0073937,FBpp0100179,FBpp0289914,FBpp0085300,FBpp0088955,FBpp0082801,FBpp0079710,FBpp0071583,FBpp0089379,FBpp0071255,FBpp0082541,FBpp0081102,FBpp0078649,FBpp0075485,FBpp0084175,FBpp0078731,FBpp0080433,FBpp0079000,FBpp0087733,FBpp0289181,FBpp0305077,FBpp0081253,FBpp0085590,FBpp0076077,FBpp0298274,FBpp0075106,FBpp0075527,FBpp0077938,FBpp0076952,FBpp0075344,FBpp0072595,FBpp0077130,FBpp0078437,FBpp0304117,FBpp0073947,FBpp0077359,FBpp0289146,FBpp0083761,FBpp0084839,FBpp0076872,FBpp0293954,FBpp0080794,FBpp0301746,FBpp0083039,FBpp0078630,FBpp0289717,FBpp0079908,FBpp0076635,FBpp0084525,FBpp0074364,FBpp0111290,FBpp0081288,FBpp0080046,FBpp0071254,FBpp0082759,FBpp0084947,FBpp0302976,FBpp0078413,FBpp0079697,FBpp0076467,FBpp0302921,FBpp0088669,FBpp0081843,FBpp0080674,FBpp0074816,FBpp0079203,FBpp0070935,FBpp0303743,FBpp0071478,FBpp0086120,FBpp0082243,FBpp0302807,FBpp0075138,FBpp0071262,FBpp0083939,FBpp0075498,FBpp0075246,FBpp0089424,FBpp0112119,FBpp0081475,FBpp0070952,FBpp0304499,FBpp0075581,FBpp0073313,FBpp0303258,FBpp0110582,FBpp0084191,FBpp0071849,FBpp0289099,FBpp0303475,FBpp0099488,FBpp0088843,FBpp0305184,FBpp0074806,FBpp0072216,FBpp0070751,FBpp0081123,FBpp0078565,FBpp0078490,FBpp0070431,FBpp0082483,FBpp0112173,FBpp0297878,FBpp0075941,FBpp0085693,FBpp0071562,FBpp0302863,FBpp0304747,FBpp0073783,FBpp0081889,FBpp0086570,FBpp0074888,FBpp0072456,FBpp0305600,FBpp0087649,FBpp0070116,FBpp0290743,FBpp0080593,FBpp0085131,FBpp0086627,FBpp0305599,FBpp0087437,FBpp0298289,FBpp0075348,FBpp0290265,FBpp0305671,FBpp0076090,FBpp0073041,FBpp0078326,FBpp0304733,FBpp0075353,FBpp0084591,FBpp0073906,FBpp0070317,FBpp0302939,FBpp0305612,FBpp0072258,FBpp0303745,FBpp0073228,FBpp0300985,FBpp0085607,FBpp0082320,FBpp0304166,FBpp0086875,FBpp0078819,FBpp0078450,FBpp0080939,FBpp0077550,FBpp0074009,FBpp0084144,FBpp0301582,FBpp0077427,FBpp0072094,FBpp0083766,FBpp0071217,FBpp0082019,FBpp0111805,FBpp0081552,FBpp0088886,FBpp0077384,FBpp0081356,FBpp0074348,FBpp0082897,FBpp0292343,FBpp0291850,FBpp0075395,FBpp0079751,FBpp0082427,FBpp0073945,FBpp0300511,FBpp0289512,FBpp0112070,FBpp0074993,FBpp0073537,FBpp0075612,FBpp0088033,FBpp0085417,FBpp0071210,FBpp0074080,FBpp0289862,FBpp0074104,FBpp0074466,FBpp0075631,FBpp0071365,FBpp0086482,FBpp0075091,FBpp0082161,FBpp0082932,FBpp0073910,FBpp0081613,FBpp0076481,FBpp0305835,FBpp0082246,FBpp0078287,FBpp0086272,FBpp0079899,FBpp0288958,FBpp0075861,FBpp0088643,FBpp0079304,FBpp0070250,FBpp0289272,FBpp0075865,FBpp0303527,FBpp0080408,FBpp0083673,FBpp0082588,FBpp0074970,FBpp0089402,FBpp0304028,FBpp0080200,FBpp0070481
cluster    2   32   43FBpp0080131,FBpp0099733,FBpp0080979,FBpp0081649,FBpp0087745,FBpp0079309,FBpp0110074,FBpp0088125,FBpp0082880,FBpp0085060,FBpp0110068,FBpp0304235,FBpp0074612,FBpp0088124,FBpp0072325,FBpp0289587,FBpp0074805,FBpp0081199,FBpp0088123,FBpp0077596,FBpp0079179,FBpp0080129,FBpp0305543,FBpp0079747,FBpp0070801,FBpp0082594,FBpp0079094,FBpp0084963,FBpp0077594,FBpp0085523,FBpp0290084,FBpp0079528
cluster    3   22   28FBpp0306147,FBpp0078721,FBpp0300982,FBpp0071140,FBpp0072036,FBpp0077451,FBpp0073073,FBpp0081581,FBpp0087164,FBpp0073135,FBpp0085703,FBpp0087739,FBpp0082834,FBpp0071844,FBpp0073426,FBpp0088273,FBpp0080802,FBpp0088881,FBpp0292397,FBpp0071536,FBpp0075401,FBpp0085176
cluster    4   20   37FBpp0082980,FBpp0085609,FBpp0072457,FBpp0070160,FBpp0081787,FBpp0112462,FBpp0084211,FBpp0078361,FBpp0076599,FBpp0074821,FBpp0087251,FBpp0070295,FBpp0081639,FBpp0305737,FBpp0079166,FBpp0289888,FBpp0074926,FBpp0078496,FBpp0112954,FBpp0077722
cluster    5   20   29FBpp0070584,FBpp0081458,FBpp0086468,FBpp0087108,FBpp0085483,FBpp0087668,FBpp0288461,FBpp0082139,FBpp0078349,FBpp0083793,FBpp0080495,FBpp0080010,FBpp0083071,FBpp0086474,FBpp0305304,FBpp0078234,FBpp0288471,FBpp0110131,FBpp0083214,FBpp0085786
cluster    6   17   22FBpp0304504,FBpp0070072,FBpp0072477,FBpp0083863,FBpp0070071,FBpp0081318,FBpp0084334,FBpp0084332,FBpp0084355,FBpp0071864,FBpp0084328,FBpp0070482,FBpp0303693,FBpp0080631,FBpp0075857,FBpp0070073,FBpp0073317
cluster    7   14   23FBpp0073338,FBpp0071593,FBpp0112504,FBpp0074536,FBpp0073606,FBpp0288960,FBpp0110179,FBpp0079894,FBpp0082080,FBpp0083163,FBpp0073033,FBpp0085338,FBpp0076949,FBpp0070474
cluster    8   14   18FBpp0078962,FBpp0289150,FBpp0082624,FBpp0082383,FBpp0088260,FBpp0080628,FBpp0077055,FBpp0290580,FBpp0288481,FBpp0289501,FBpp0076386,FBpp0077218,FBpp0085279,FBpp0072679
cluster    9   14   24FBpp0072691,FBpp0300826,FBpp0084340,FBpp0077896,FBpp0074859,FBpp0087704,FBpp0078598,FBpp0076658,FBpp0088666,FBpp0305334,FBpp0074278,FBpp0081310,FBpp0297275,FBpp0076869
cluster   10   12   15FBpp0290701,FBpp0304954,FBpp0071585,FBpp0302578,FBpp0087527,FBpp0303214,FBpp0081882,FBpp0072215,FBpp0070154,FBpp0078822,FBpp0304268,FBpp0076858
cluster   11   10    9FBpp0089115,FBpp0071007,FBpp0070105,FBpp0077189,FBpp0077188,FBpp0084352,FBpp0078726,FBpp0087197,FBpp0074234,FBpp0071809
cluster   12   10   10FBpp0084964,FBpp0072579,FBpp0080683,FBpp0082792,FBpp0303868,FBpp0070722,FBpp0289986,FBpp0301562,FBpp0077250,FBpp0077928
cluster   13   10   11FBpp0076956,FBpp0079324,FBpp0071279,FBpp0079233,FBpp0304680,FBpp0304506,FBpp0088527,FBpp0080994,FBpp0081362,FBpp0075946
cluster   14    9    9FBpp0072780,FBpp0072761,FBpp0271837,FBpp0078236,FBpp0289343,FBpp0070196,FBpp0289676,FBpp0293660,FBpp0076590
cluster   15    9   29FBpp0079992,FBpp0082787,FBpp0087095,FBpp0087764,FBpp0071226,FBpp0081401,FBpp0073902,FBpp0083683,FBpp0075794
cluster   16    9    9FBpp0082112,FBpp0083954,FBpp0289500,FBpp0304299,FBpp0080292,FBpp0302594,FBpp0072861,FBpp0078390,FBpp0072099
cluster   17    9    8FBpp0082356,FBpp0082542,FBpp0087757,FBpp0305939,FBpp0072825,FBpp0084240,FBpp0072334,FBpp0073965,FBpp0079914
cluster   18    9   14FBpp0074829,FBpp0290948,FBpp0075932,FBpp0075689,FBpp0075998,FBpp0289374,FBpp0077417,FBpp0110178,FBpp0072449
cluster   19    8    7FBpp0071136,FBpp0084549,FBpp0080080,FBpp0305215,FBpp0074051,FBpp0081569,FBpp0271525,FBpp0296952
cluster   20    8    9FBpp0082819,FBpp0079711,FBpp0081262,FBpp0078227,FBpp0080679,FBpp0084172,FBpp0086399,FBpp0071818
cluster   21    8    7FBpp0073940,FBpp0082793,FBpp0082829,FBpp0082826,FBpp0081161,FBpp0081138,FBpp0081732,FBpp0089393
cluster   22    8    8FBpp0079448,FBpp0291777,FBpp0087524,FBpp0075283,FBpp0290718,FBpp0088350,FBpp0085089,FBpp0084919
cluster   23    7    9FBpp0084133,FBpp0073441,FBpp0070643,FBpp0081576,FBpp0290661,FBpp0071663,FBpp0080608
cluster   24    7    6FBpp0072101,FBpp0080661,FBpp0074370,FBpp0073014,FBpp0301569,FBpp0083032,FBpp0085394
cluster   25    7    7FBpp0074756,FBpp0303820,FBpp0085423,FBpp0072164,FBpp0081704,FBpp0074910,FBpp0290468
cluster   26    7    7FBpp0084783,FBpp0271922,FBpp0079869,FBpp0304416,FBpp0305385,FBpp0072571,FBpp0087059
cluster   27    7    6FBpp0074638,FBpp0289203,FBpp0075915,FBpp0304756,FBpp0111921,FBpp0290308,FBpp0076857
cluster   28    6    5FBpp0077802,FBpp0070169,FBpp0076478,FBpp0111428,FBpp0085455,FBpp0075046
cluster   29    6    6FBpp0083028,FBpp0071241,FBpp0290236,FBpp0075096,FBpp0073378,FBpp0086329
cluster   30    6    5FBpp0305272,FBpp0300338,FBpp0072200,FBpp0086326,FBpp0074517,FBpp0112201
cluster   31    6    5FBpp0289244,FBpp0110250,FBpp0084716,FBpp0298271,FBpp0072491,FBpp0081202
cluster   32    6    6FBpp0080044,FBpp0080284,FBpp0086996,FBpp0084043,FBpp0073806,FBpp0076338
cluster   33    6    6FBpp0297826,FBpp0080999,FBpp0074786,FBpp0078338,FBpp0080790,FBpp0077872
cluster   34    6    5FBpp0080448,FBpp0086590,FBpp0075999,FBpp0110523,FBpp0077868,FBpp0078425
cluster   35    6    5FBpp0076026,FBpp0087262,FBpp0112105,FBpp0088122,FBpp0076712,FBpp0082351
cluster   36    6    5FBpp0291385,FBpp0086244,FBpp0111987,FBpp0078104,FBpp0082068,FBpp0079217
cluster   37    6    5FBpp0084507,FBpp0111746,FBpp0082539,FBpp0084431,FBpp0304588,FBpp0083832
cluster   38    5    8FBpp0071123,FBpp0070333,FBpp0070795,FBpp0079294,FBpp0088455
cluster   39    5    4FBpp0086375,FBpp0070672,FBpp0112072,FBpp0085122,FBpp0073975
cluster   40    5    6FBpp0083549,FBpp0081328,FBpp0079352,FBpp0077173,FBpp0071024
cluster   41    5    6FBpp0072135,FBpp0079472,FBpp0072560,FBpp0071095,FBpp0079454
cluster   42    5    4FBpp0071302,FBpp0074907,FBpp0075318,FBpp0305104,FBpp0075168
cluster   43    5    5FBpp0071476,FBpp0084110,FBpp0099887,FBpp0086074,FBpp0074386
cluster   44    5    4FBpp0080901,FBpp0291175,FBpp0290582,FBpp0072496,FBpp0087930
cluster   45    5    5FBpp0084842,FBpp0082713,FBpp0077627,FBpp0073007,FBpp0075250
cluster   46    5    4FBpp0074562,FBpp0077962,FBpp0081336,FBpp0073066,FBpp0080089
cluster   47    5    5FBpp0079502,FBpp0289821,FBpp0288426,FBpp0289395,FBpp0073155
cluster   48    5    4FBpp0074817,FBpp0082798,FBpp0076237,FBpp0087788,FBpp0073358
cluster   49    5    4FBpp0077149,FBpp0084411,FBpp0073525,FBpp0080011,FBpp0290362
cluster   50    5    4FBpp0304792,FBpp0082535,FBpp0073955,FBpp0113106,FBpp0077121
cluster   51    5    5FBpp0079755,FBpp0087092,FBpp0079754,FBpp0088574,FBpp0074842
cluster   52    5    5FBpp0086568,FBpp0074940,FBpp0082908,FBpp0085803,FBpp0086518
cluster   53    5    5FBpp0076224,FBpp0301282,FBpp0076182,FBpp0076225,FBpp0100010
cluster   54    5    4FBpp0088200,FBpp0076350,FBpp0080687,FBpp0088864,FBpp0087144
cluster   55    5    4FBpp0300931,FBpp0085657,FBpp0076547,FBpp0080675,FBpp0297182
cluster   56    5    4FBpp0081101,FBpp0081086,FBpp0291365,FBpp0305435,FBpp0300825
cluster   57    5    7FBpp0086042,FBpp0081279,FBpp0289977,FBpp0084347,FBpp0298004
cluster   58    4    3FBpp0077129,FBpp0070308,FBpp0078729,FBpp0087307
cluster   59    4    3FBpp0076340,FBpp0070717,FBpp0074608,FBpp0075447
cluster   60    4    4FBpp0070742,FBpp0086336,FBpp0292257,FBpp0085560
cluster   61    4    3FBpp0289508,FBpp0079919,FBpp0070768,FBpp0073057
cluster   62    4    3FBpp0288463,FBpp0271880,FBpp0082657,FBpp0071014
cluster   63    4    6FBpp0071739,FBpp0298043,FBpp0289907,FBpp0071754
cluster   64    4    3FBpp0072277,FBpp0293785,FBpp0079183,FBpp0290823
cluster   65    4    3FBpp0086226,FBpp0100079,FBpp0080148,FBpp0072852
cluster   66    4    3FBpp0072878,FBpp0079267,FBpp0072880,FBpp0072877
cluster   67    4    3FBpp0073010,FBpp0076840,FBpp0074419,FBpp0291001
cluster   68    4    3FBpp0073526,FBpp0297991,FBpp0074105,FBpp0082953
cluster   69    4    3FBpp0086971,FBpp0086895,FBpp0077736,FBpp0073731
cluster   70    4    3FBpp0073917,FBpp0079855,FBpp0088673,FBpp0083054
cluster   71    4    3FBpp0079162,FBpp0074003,FBpp0087097,FBpp0305307
cluster   72    4    3FBpp0083749,FBpp0304253,FBpp0074523,FBpp0083799
cluster   73    4    3FBpp0079623,FBpp0304827,FBpp0074544,FBpp0076535
cluster   74    4    3FBpp0078893,FBpp0083227,FBpp0074843,FBpp0076520
cluster   75    4    5FBpp0076326,FBpp0298351,FBpp0082600,FBpp0079875
cluster   76    4    4FBpp0076621,FBpp0077718,FBpp0087297,FBpp0085718
cluster   77    4    3FBpp0076818,FBpp0086083,FBpp0086085,FBpp0086069
cluster   78    4    3FBpp0300515,FBpp0292041,FBpp0077399,FBpp0086515
cluster   79    4    3FBpp0081847,FBpp0086957,FBpp0292109,FBpp0077878
cluster   80    4    3FBpp0081866,FBpp0085077,FBpp0079490,FBpp0081561
cluster   81    4    3FBpp0083238,FBpp0304758,FBpp0080033,FBpp0080059
cluster   82    4    3FBpp0081153,FBpp0085721,FBpp0085327,FBpp0086328
cluster   83    3    2FBpp0070057,FBpp0080551,FBpp0083050
cluster   84    3    2FBpp0083463,FBpp0070070,FBpp0305260
cluster   85    3    3FBpp0300161,FBpp0074784,FBpp0070087
cluster   86    3    2FBpp0070165,FBpp0080860,FBpp0087317
cluster   87    3    2FBpp0070552,FBpp0076101,FBpp0070390
cluster   88    3    2FBpp0084140,FBpp0070476,FBpp0289092
cluster   89    3    2FBpp0078157,FBpp0078810,FBpp0070899
cluster   90    3    3FBpp0073568,FBpp0070924,FBpp0087999
cluster   91    3    2FBpp0073239,FBpp0071435,FBpp0071021
cluster   92    3    2FBpp0071192,FBpp0081961,FBpp0079549
cluster   93    3    2FBpp0071461,FBpp0077549,FBpp0302572
cluster   94    3    2FBpp0292003,FBpp0073296,FBpp0071501
cluster   95    3    2FBpp0087860,FBpp0071618,FBpp0305959
cluster   96    3    2FBpp0071669,FBpp0078156,FBpp0303775
cluster   97    3    2FBpp0071779,FBpp0288419,FBpp0084614
cluster   98    3    2FBpp0072003,FBpp0077355,FBpp0072168
cluster   99    3    2FBpp0111775,FBpp0080707,FBpp0072054
cluster  100    3    2FBpp0072523,FBpp0075683,FBpp0298292
cluster  101    3    3FBpp0076329,FBpp0072626,FBpp0305188
cluster  102    3    2FBpp0072643,FBpp0303586,FBpp0076971
cluster  103    3    2FBpp0081547,FBpp0303964,FBpp0073293
cluster  104    3    2FBpp0292197,FBpp0073500,FBpp0082599
cluster  105    3    3FBpp0077750,FBpp0073640,FBpp0086057
cluster  106    3    2FBpp0076204,FBpp0111839,FBpp0073678
cluster  107    3    2FBpp0085107,FBpp0086555,FBpp0074684
cluster  108    3    2FBpp0074714,FBpp0085870,FBpp0087978
cluster  109    3    2FBpp0074785,FBpp0291035,FBpp0303995
cluster  110    3    2FBpp0083943,FBpp0079251,FBpp0074824
cluster  111    3    2FBpp0081139,FBpp0074853,FBpp0079882
cluster  112    3    2FBpp0110403,FBpp0082276,FBpp0075117
cluster  113    3    2FBpp0075748,FBpp0288501,FBpp0291628
cluster  114    3    2FBpp0076348,FBpp0305731,FBpp0083838
cluster  115    3    2FBpp0080746,FBpp0078144,FBpp0083515
cluster  116    3    2FBpp0078342,FBpp0083931,FBpp0085842
cluster  117    3    2FBpp0078964,FBpp0087941,FBpp0089177
cluster  118    3    2FBpp0085075,FBpp0087972,FBpp0079207
cluster  119    3    2FBpp0079219,FBpp0086773,FBpp0083917
cluster  120    3    2FBpp0084040,FBpp0079701,FBpp0305197
cluster  121    3    2FBpp0080015,FBpp0084124,FBpp0083200
cluster  122    3    2FBpp0080297,FBpp0086842,FBpp0111402
cluster  123    3    2FBpp0081088,FBpp0087766,FBpp0084912
cluster  124    3    3FBpp0081282,FBpp0082644,FBpp0289882
cluster  125    3    2FBpp0085413,FBpp0081347,FBpp0085134
cluster  126    3    2FBpp0084383,FBpp0301834,FBpp0081609
cluster  127    3    2FBpp0292856,FBpp0289101,FBpp0082647
cluster  128    3    2FBpp0301546,FBpp0301600,FBpp0082709
cluster  129    3    2FBpp0289058,FBpp0085878,FBpp0082816
cluster  130    3    2FBpp0083969,FBpp0083132,FBpp0088773
cluster  131    3    2FBpp0083213,FBpp0083959,FBpp0087806
cluster  132    3    2FBpp0303926,FBpp0083431,FBpp0305929
cluster  133    3    2FBpp0083457,FBpp0087266,FBpp0086289
cluster  134    3    2FBpp0083574,FBpp0113052,FBpp0085904
cluster  135    3    2FBpp0083704,FBpp0304695,FBpp0084418
cluster  136    3    3FBpp0085115,FBpp0084819,FBpp0110262
cluster  137    3    2FBpp0086579,FBpp0086017,FBpp0088870
cluster  138    3    2FBpp0301994,FBpp0288697,FBpp0086898
cluster  139    3    2FBpp0292059,FBpp0099910,FBpp0289718
cluster  140    3    2FBpp0111766,FBpp0290370,FBpp0291978
cluster  141    3    2FBpp0271806,FBpp0289521,FBpp0111858
cluster  142    2    1FBpp0070074,FBpp0087879
cluster  143    2    1FBpp0076250,FBpp0070083
cluster  144    2    1FBpp0075595,FBpp0070181
cluster  145    2    1FBpp0073209,FBpp0070204
cluster  146    2    1FBpp0070221,FBpp0081367
cluster  147    2    1FBpp0071431,FBpp0070280
cluster  148    2    1FBpp0070335,FBpp0079708
cluster  149    2    1FBpp0079675,FBpp0070350
cluster  150    2    1FBpp0079907,FBpp0070419
cluster  151    2    1FBpp0070545,FBpp0074205
cluster  152    2    1FBpp0070589,FBpp0076440
cluster  153    2    1FBpp0070622,FBpp0075816
cluster  154    2    1FBpp0075572,FBpp0070744
cluster  155    2    1FBpp0070757,FBpp0075325
cluster  156    2    1FBpp0070748,FBpp0099986
cluster  157    2    1FBpp0070775,FBpp0082060
cluster  158    2    1FBpp0070792,FBpp0077198
cluster  159    2    1FBpp0302657,FBpp0070832
cluster  160    2    1FBpp0079595,FBpp0070931
cluster  161    2    1FBpp0083990,FBpp0070963
cluster  162    2    1FBpp0083729,FBpp0071063
cluster  163    2    1FBpp0071832,FBpp0071104
cluster  164    2    1FBpp0071138,FBpp0084021
cluster  165    2    1FBpp0082979,FBpp0071173
cluster  166    2    1FBpp0071235,FBpp0073572
cluster  167    2    1FBpp0075209,FBpp0071307
cluster  168    2    1FBpp0071450,FBpp0080100
cluster  169    2    1FBpp0071462,FBpp0082412
cluster  170    2    1FBpp0081402,FBpp0071502
cluster  171    2    1FBpp0071597,FBpp0112286
cluster  172    2    1FBpp0084012,FBpp0071745
cluster  173    2    1FBpp0087722,FBpp0071894
cluster  174    2    1FBpp0085699,FBpp0071950
cluster  175    2    1FBpp0076243,FBpp0071938
cluster  176    2    1FBpp0071969,FBpp0085809
cluster  177    2    1FBpp0071971,FBpp0076076
cluster  178    2    1FBpp0072070,FBpp0080563
cluster  179    2    1FBpp0072269,FBpp0085761
cluster  180    2    1FBpp0083456,FBpp0072331
cluster  181    2    1FBpp0088363,FBpp0072365
cluster  182    2    1FBpp0072472,FBpp0086139
cluster  183    2    1FBpp0072701,FBpp0099770
cluster  184    2    1FBpp0306019,FBpp0072703
cluster  185    2    1FBpp0072782,FBpp0271715
cluster  186    2    1FBpp0079321,FBpp0072820
cluster  187    2    1FBpp0113095,FBpp0072848
cluster  188    2    1FBpp0086939,FBpp0072944
cluster  189    2    1FBpp0110117,FBpp0073050
cluster  190    2    1FBpp0073069,FBpp0089220
cluster  191    2    1FBpp0073248,FBpp0288875
cluster  192    2    1FBpp0081185,FBpp0073355
cluster  193    2    1FBpp0305727,FBpp0073360
cluster  194    2    1FBpp0079411,FBpp0073457
cluster  195    2    1FBpp0083967,FBpp0073551
cluster  196    2    1FBpp0073578,FBpp0291414
cluster  197    2    1FBpp0073669,FBpp0085251
cluster  198    2    1FBpp0076666,FBpp0073725
cluster  199    2    1FBpp0073761,FBpp0303762
cluster  200    2    1FBpp0081591,FBpp0073816
cluster  201    2    1FBpp0089324,FBpp0073792
cluster  202    2    1FBpp0073853,FBpp0084782
cluster  203    2    1FBpp0292515,FBpp0073891
cluster  204    2    1FBpp0078469,FBpp0074025
cluster  205    2    1FBpp0074103,FBpp0074133
cluster  206    2    1FBpp0074511,FBpp0081246
cluster  207    2    1FBpp0074516,FBpp0078372
cluster  208    2    1FBpp0074564,FBpp0088213
cluster  209    2    1FBpp0081723,FBpp0074693
cluster  210    2    1FBpp0074807,FBpp0087625
cluster  211    2    1FBpp0074813,FBpp0294018
cluster  212    2    1FBpp0292147,FBpp0074992
cluster  213    2    1FBpp0305956,FBpp0074991
cluster  214    2    1FBpp0075156,FBpp0302002
cluster  215    2    1FBpp0075148,FBpp0076652
cluster  216    2    1FBpp0289215,FBpp0075692
cluster  217    2    1FBpp0075721,FBpp0082386
cluster  218    2    1FBpp0088615,FBpp0075732
cluster  219    2    1FBpp0289061,FBpp0075750
cluster  220    2    1FBpp0075762,FBpp0076861
cluster  221    2    1FBpp0075826,FBpp0075891
cluster  222    2    1FBpp0075912,FBpp0306233
cluster  223    2    1FBpp0075928,FBpp0080523
cluster  224    2    1FBpp0075984,FBpp0076921
cluster  225    2    1FBpp0076083,FBpp0083025
cluster  226    2    1FBpp0076145,FBpp0291761
cluster  227    2    1FBpp0113069,FBpp0076164
cluster  228    2    1FBpp0088492,FBpp0076167
cluster  229    2    1FBpp0076448,FBpp0088428
cluster  230    2    1FBpp0290511,FBpp0076550
cluster  231    2    1FBpp0076608,FBpp0305294
cluster  232    2    1FBpp0079313,FBpp0076790
cluster  233    2    1FBpp0087056,FBpp0076942
cluster  234    2    1FBpp0078268,FBpp0077026
cluster  235    2    1FBpp0086073,FBpp0077134
cluster  236    2    1FBpp0077144,FBpp0084691
cluster  237    2    1FBpp0077400,FBpp0291079
cluster  238    2    1FBpp0083029,FBpp0077444
cluster  239    2    1FBpp0303850,FBpp0077626
cluster  240    2    1FBpp0078362,FBpp0077676
cluster  241    2    1FBpp0088783,FBpp0077850
cluster  242    2    1FBpp0077841,FBpp0079317
cluster  243    2    1FBpp0077874,FBpp0099837
cluster  244    2    1FBpp0079574,FBpp0077934
cluster  245    2    1FBpp0099514,FBpp0077941
cluster  246    2    1FBpp0077950,FBpp0294001
cluster  247    2    1FBpp0297995,FBpp0078304
cluster  248    2    1FBpp0085418,FBpp0078331
cluster  249    2    1FBpp0078337,FBpp0302877
cluster  250    2    1FBpp0302051,FBpp0078424
cluster  251    2    1FBpp0086475,FBpp0078510
cluster  252    2    1FBpp0078512,FBpp0289682
cluster  253    2    1FBpp0083150,FBpp0078693
cluster  254    2    1FBpp0081028,FBpp0078759
cluster  255    2    1FBpp0078842,FBpp0083604
cluster  256    2    1FBpp0078813,FBpp0083626
cluster  257    2    1FBpp0078854,FBpp0086114
cluster  258    2    1FBpp0079066,FBpp0083775
cluster  259    2    1FBpp0110478,FBpp0079091
cluster  260    2    1FBpp0079308,FBpp0086868
cluster  261    2    1FBpp0079318,FBpp0304316
cluster  262    2    1FBpp0086253,FBpp0079361
cluster  263    2    1FBpp0079371,FBpp0087587
cluster  264    2    1FBpp0084120,FBpp0079462
cluster  265    2    1FBpp0079547,FBpp0082349
cluster  266    2    1FBpp0080407,FBpp0079578
cluster  267    2    1FBpp0085511,FBpp0079596
cluster  268    2    1FBpp0087152,FBpp0079648
cluster  269    2    1FBpp0079732,FBpp0088692
cluster  270    2    1FBpp0080031,FBpp0084027
cluster  271    2    1FBpp0087086,FBpp0080057
cluster  272    2    1FBpp0080134,FBpp0080135
cluster  273    2    1FBpp0082026,FBpp0080156
cluster  274    2    1FBpp0083529,FBpp0080207
cluster  275    2    1FBpp0080212,FBpp0084758
cluster  276    2    1FBpp0080261,FBpp0089410
cluster  277    2    1FBpp0081861,FBpp0080322
cluster  278    2    1FBpp0080513,FBpp0083137
cluster  279    2    1FBpp0290849,FBpp0080537
cluster  280    2    1FBpp0080547,FBpp0088054
cluster  281    2    1FBpp0080554,FBpp0080688
cluster  282    2    1FBpp0289422,FBpp0080582
cluster  283    2    1FBpp0080783,FBpp0080781
cluster  284    2    1FBpp0099388,FBpp0080817
cluster  285    2    1FBpp0304913,FBpp0080859
cluster  286    2    1FBpp0111291,FBpp0081087
cluster  287    2    1FBpp0081206,FBpp0292842
cluster  288    2    1FBpp0081457,FBpp0081464
cluster  289    2    1FBpp0081476,FBpp0081741
cluster  290    2    1FBpp0081483,FBpp0082178
cluster  291    2    1FBpp0301157,FBpp0081525
cluster  292    2    1FBpp0290859,FBpp0081629
cluster  293    2    1FBpp0084917,FBpp0081722
cluster  294    2    1FBpp0083198,FBpp0081946
cluster  295    2    1FBpp0084873,FBpp0081962
cluster  296    2    1FBpp0086370,FBpp0081993
cluster  297    2    1FBpp0305489,FBpp0082066
cluster  298    2    1FBpp0110105,FBpp0082156
cluster  299    2    1FBpp0086304,FBpp0082359
cluster  300    2    1FBpp0082399,FBpp0112409
cluster  301    2    1FBpp0088079,FBpp0082538
cluster  302    2    1FBpp0082549,FBpp0087398
cluster  303    2    1FBpp0083774,FBpp0082625
cluster  304    2    1FBpp0112084,FBpp0082720
cluster  305    2    1FBpp0082812,FBpp0290949
cluster  306    2    1FBpp0082895,FBpp0083127
cluster  307    2    1FBpp0082916,FBpp0084312
cluster  308    2    1FBpp0082981,FBpp0289204
cluster  309    2    1FBpp0083045,FBpp0087515
cluster  310    2    1FBpp0083088,FBpp0302616
cluster  311    2    1FBpp0304952,FBpp0083114
cluster  312    2    1FBpp0083352,FBpp0087869
cluster  313    2    1FBpp0291478,FBpp0083356
cluster  314    2    1FBpp0083432,FBpp0300435
cluster  315    2    1FBpp0083533,FBpp0087774
cluster  316    2    1FBpp0293006,FBpp0083726
cluster  317    2    1FBpp0083731,FBpp0084938
cluster  318    2    1FBpp0087819,FBpp0083995
cluster  319    2    1FBpp0301783,FBpp0084416
cluster  320    2    1FBpp0084396,FBpp0300789
cluster  321    2    1FBpp0100150,FBpp0084420
cluster  322    2    1FBpp0084505,FBpp0084506
cluster  323    2    1FBpp0084440,FBpp0087974
cluster  324    2    1FBpp0084528,FBpp0305896
cluster  325    2    1FBpp0084540,FBpp0084615
cluster  326    2    1FBpp0084543,FBpp0290673
cluster  327    2    1FBpp0084646,FBpp0084647
cluster  328    2    1FBpp0084801,FBpp0293581
cluster  329    2    1FBpp0084927,FBpp0110277
cluster  330    2    1FBpp0085155,FBpp0304160
cluster  331    2    1FBpp0085643,FBpp0085633
cluster  332    2    1FBpp0288951,FBpp0085662
cluster  333    2    1FBpp0085706,FBpp0087440
cluster  334    2    1FBpp0099678,FBpp0085804
cluster  335    2    1FBpp0085834,FBpp0088658
cluster  336    2    1FBpp0086020,FBpp0290276
cluster  337    2    1FBpp0305298,FBpp0086056
cluster  338    2    1FBpp0086119,FBpp0291587
cluster  339    2    1FBpp0300234,FBpp0086405
cluster  340    2    1FBpp0305858,FBpp0086498
cluster  341    2    1FBpp0086485,FBpp0303248
cluster  342    2    1FBpp0111783,FBpp0086614
cluster  343    2    1FBpp0087714,FBpp0086803
cluster  344    2    1FBpp0111404,FBpp0086930
cluster  345    2    1FBpp0293886,FBpp0086903
cluster  346    2    1FBpp0086994,FBpp0306143
cluster  347    2    1FBpp0087122,FBpp0304242
cluster  348    2    1FBpp0303859,FBpp0087120
cluster  349    2    1FBpp0087424,FBpp0297743
cluster  350    2    1FBpp0088585,FBpp0087483
cluster  351    2    1FBpp0304599,FBpp0087493
cluster  352    2    1FBpp0087559,FBpp0292065
cluster  353    2    1FBpp0088181,FBpp0297720
cluster  354    2    1FBpp0088162,FBpp0089409
cluster  355    2    1FBpp0113067,FBpp0088599
cluster  356    2    1FBpp0088679,FBpp0110092
cluster  357    2    1FBpp0110147,FBpp0099486
cluster  358    2    1FBpp0305308,FBpp0099982
cluster  359    2    1FBpp0305367,FBpp0099952
cluster  360    2    1FBpp0297080,FBpp0111412
cluster  361    2    1FBpp0289419,FBpp0111760
cluster  362    2    1FBpp0111689,FBpp0289201
cluster  363    2    1FBpp0112499,FBpp0293083
cluster  364    2    1FBpp0289746,FBpp0291104
cluster  365    2    1FBpp0290783,FBpp0297903
cluster  366    2    1FBpp0305547,FBpp0291443
cluster  367    2    1FBpp0291496,FBpp0291497
cluster  368    2    1FBpp0291631,FBpp0305937
cluster  369    2    1FBpp0303905,FBpp0300445
cluster  370    2    1FBpp0304074,FBpp0300637
cluster  371    2    1FBpp0302954,FBpp0303126
cluster  372    2    1FBpp0304478,FBpp0304479
